# Performing Reflexive Toxin A/B Enzyme Immunoassay in a Two-Step Algorithm for the Diagnosis of *Clostridium difficile* Infection Has Limited Clinical Utility

**DOI:** 10.1101/384214

**Authors:** M Mounajjed, R Pease, B Jung-Hynes, N Safdar, D Chen

## Abstract

The differentiation of *Clostridium difficile* infection (CDI) from colonization is challenged by the suboptimal clinical specificity of nucleic acid amplification tests (NAAT). In this study, we examined the utility of testing for toxin via enzyme immunoassay (EIA) in specimens already tested by NAAT for the diagnosis of CDI in an attempt to differentiate colonization from infection. We tested 59 stool samples for the presence of *C. difficile* toxin B gene by NAAT followed by EIAs for glutamate dehydrogenase (GDH EIA) and toxins A and B (Toxin EIA). Two infectious disease physicians independently reviewed the patients’ electronic medical records retrospectively to categorize each patient as CDI-Likely, CDI-Unlikely, or CDI-Indeterminate. Clinical sensitivities and specificities were calculated using 3 definitions of “true” CDI status, being: (1) concordance between both reviewers, (2) concordance, and CDI-Indeterminate/discordant cases classified as CDI-Likely, and (3) concordance, and CDI-Indeterminate/discordant cases classified as CDI-Unlikely. Based on these definitions, clinical sensitivity and specificity for NAAT was 100% and 49-94%, GDH EIA was 83-85% and 43-89%, and Toxin EIA was 39-42% and 83-100%, respectively. 85% (22 of 26) of patients who were NAAT-positive but Toxin EIA-negative symptomatically benefited from treatment for CDI. The addition of EIA to NAAT for CDI diagnosis had limited utility for differentiating colonization from CDI and could have led to under treatment of patients with CDI.

## INTRODUCTION

*Clostridioides* (formerly *Clostridium*) *difficile* is the most commonly recognized cause of infectious diarrhea in healthcare settings, and is the most commonly reported pathogen among healthcare-associated infections of adults in the United States (1). However, asymptomatic colonization has been found in acute care hospitals at an incidence of 3%-26% in adult inpatients (2-4). Differentiating patients who are colonized from those who have active CDI remains a diagnostic challenge. While NAAT has a high clinical sensitivity, its clinical specificity is suboptimal for diagnosing patients who have CDI(5). An array of algorithms and protocols have been created to improve diagnostic accuracy of CDI.

The 2017 Infectious Disease Society of America (IDSA) and Society for Healthcare Epidemiology of America (SHEA) guidelines for *C. difficile* recommend the use of a stool toxin test as a part of a multistep algorithm (i.e. GDH plus toxin; GDH plus toxin, arbitrated by NAAT; or NAAT plus toxin) at institutions without a formal process to submit stool specimens from only patients with unexplained new onset diarrhea with 3 or more unformed stools in a 24 hour period. At institutions that do have a policy to only submit stool for *C. difficile* testing in patients with 3 or more unformed stools in a 24-hour period, testing by NAAT alone is provided as an alternative diagnostic option (6).

Accurate diagnosis of CDI is essential. A false positive test result would lead to unnecessary antibiotic exposure and increases in costs of care, while missing a diagnosis of CDI would likely lead to increased morbidity and mortality. We undertook this study to evaluate the impact of adding toxin testing to NAAT to differentiate colonization from CDI (7).

## METHODS

This study was performed at a 500-bed, tertiary care, academic medical center in the Midwestern United States. As part of routine diagnostic workup for CDI, unformed stool specimens were submitted to the clinical microbiology laboratory to detect the presence of *C. difficile* toxin B gene via the Xpert *C. difficile*/Epi NAAT (Cepheid, Sunnyvale, CA). For this study, unformed stool samples, nonselectively chosen based on investigator availability from January to May 2017, were subsequently tested with the TECHLAB C. DIFF QUIK CHEK COMPLETE EIA (TECHLAB, Blacksburg, VA) according to manufacturer instructions to evaluate for the presence of GDH, and toxins A and B.

Two infectious disease physicians independently reviewed electronic medical records of the corresponding patients. Reviewers were blinded to the EIA results and to each other’s assessment. Blinding for NAAT results was not possible since the results could be readily seen within the patients’ electronic medical records. The following criteria adapted and modified from the 2010 IDSA guidelines were utilized by the reviewers in assessing disease status: presence of diarrhea defined as 3 or more unformed stools within a 24-hour period, antibiotic use within 90 days of symptom onset, prior history of CDI, improvement of symptoms with CDI treatment, and colonoscopy findings, if performed (8, 9). Since there is no gold standard for diagnosing CDI, each reviewer used the constellation of above factors, along with their clinical judgment to group patients into one of three categories: (1) CDI-Likely, (2) CDI-Unlikely, and (3) CDI-Indeterminate. Clinical sensitivity and specificity were calculated using 3 definitions of “true” CDI status: [1] both reviewers were concordant in assigning CDI-Likely or CDI-Unlikely to a patient (CDI-Indeterminate results were excluded in this calculation); [2] both reviewers were concordant in assigning CDI-Likely or CDI-Unlikely, and remaining patients (CDI-Indeterminate and discordant categorization) were included in the CDI-Likely group; and [3] both reviewers were concordant in assigning CDI-Likely or CDI-Unlikely, and remaining patients (CDI-Indeterminate and discordant categorization) were included in the CDI-Unlikely group.

## RESULTS

59 patients were included in this study. Independent categorization of CDI status by two infectious disease specialists is shown in **Table 1**. Overall percent agreement between the reviewers was 78% (46 of 59 concordant), and Kappa was 0.64 (95% confidence interval 0.47-0.80). Notably, Reviewer 1 categorized more patients as CDI-Likely (n=35) than Reviewer 2 (n=26), while Reviewer 2 categorized more patients as CDI-Indeterminate (n=12) than Reviewer 1 (n=4). Both reviewers had a similar number of CDI-Unlikely patients (n=20 and n=21). Patient age and gender were similarly distributed among each CDI status group; median age was 60 years (ranging 1 to 97 years) and there were 58% (n=34) males **(Table 2)**. NAAT was positive in 71% (42 of 59) of patients, GDH EIA in 66% (37 of 56), and Toxin EIA in 27% (16 of 59); the results of 3 GDH EIA tests could not be retrieved **(Table 3)**. Of CDI-Likely patients, 58% (14 of 24) were Toxin EIA-negative. Of NAAT-positive specimens, 62% (26 of 42) were Toxin EIA-negative, and 85% (22 of 26) of these patients symptomatically benefited from treatment for CDI. There were no (0%) instances where GDH EIA and/or Toxin EIA was positive and NAAT was negative. Using 3 different definitions of “true” CDI status, clinical sensitivity and specificity for NAAT was 100% and 49-94%, GDH EIA was 83-85% and 43-89%, and Toxin EIA was 39-42% and 83-100%, respectively **(Table 4)**.

**Table 1.** Retrospective categorization of patients with diarrhea as CDI-Likely, CDI-Unlikely, or CDI-Indeterminate based on clinical parameters by two infectious disease physicians. Percent agreement 78%, Kappa = 0.64 (95% confidence interval: 0.47-0.80).

|  |  | Reviewer 1 |  |  | Total |
| --- | --- | --- | --- | --- | --- |
| CDI Category |  | CDI-Likely | CDI-Unlikely | CDI-Indeterminate |  |
| Reviewer 2 | CDI-Likely | 24 | 2 | 0 | 26 |
|  | CDI-Unlikely | 3 | 18 | 0 | 21 |
|  | CDI-Indeterminate | 8 | 0 | 4 | 12 |
|  | Total | 35 | 20 | 4 | 59 |

**Table 2.** Age and gender of study patients and distribution by CDI category.

|  | CDI Category |  |  | Overall<br>(n=59) |
| --- | --- | --- | --- | --- |
|  | CDI-Likely<br>(n=24) | CDI-Unlikely<br>(n=18) | CDI-Indeterminate<br>or Discordant<br>(n=17) |  |
| Number Male (%) | 13 (54.2) | 11 (61.1) | 10 (58.8) | 34 (57.6) |
| Number Female (%) | 11 (45.8) | 7 (38.9) | 7 (41.2) | 25 (42.4) |
| Median Age (std dev)<br>in Years | 62.5 (19.8) | 61.0 (21.0) | 60.0 (21.9) | 61.0 (20.5) |

**Table 3:** NAAT, GDH EIA, and Toxin EIA results for patients with diarrhea distributed by CDI category.

| CDI Category | NAAT |  |  | GDH EIA |  |  |  | Toxin EIA |  |  |
| --- | --- | --- | --- | --- | --- | --- | --- | --- | --- | --- |
|  | Positive | Negative | Total | Positive | Negative | Unknown | Total | Positive | Negative | Total |
| CDI-Likely | 24 | 0 | 24 | 20 | 4 | 0 | 24 | 10 | 14 | 24 |
| CDI-Unlikely | 1 | 17 | 18 | 2 | 13 | 3* | 18 | 0 | 18 | 18 |
| CDI-Indeterminate or Discordant | 17 | 0 | 17 | 15 | 2 | 0 | 17 | 6 | 11 | 17 |
| Total | 42 | 17 | 59 | 37 | 19 | 3 | 59 | 16 | 43 | 59 |

\* GDH EIA results for 3 patients were unable to be retrieved.

**Table 4.** Clinical sensitivity and clinical specificity of NAAT, GDH EIA, and Toxin EIA calculated using 3 different definitions of “true” CDI status.

| Test | Definition of "True" CDI Status |  |  |  |  |  |
| --- | --- | --- | --- | --- | --- | --- |
|  | Concordance Between Both Reviewers in Categorizing CDI-Likely and CDI-Unlikely |  | Concordance, and CDI-Indeterminate/Discordant Considered CDI-Likely |  | Concordance, and CDI-Indeterminate/Discordant Considered CDI-Unlikely |  |
|  | Clinical Sensitivity | Clinical Specificity | Clinical Sensitivity | Clinical Specificity | Clinical Sensitivity | Clinical Specificity |
| NAAT | 100%<br>(24 of 24) | 94%<br>(17 of 18) | 100%<br>(41 of 41) | 94%<br>(17 of 18) | 100%<br>(24 of 24) | 49%<br>(17 of 35) |
| GDH EIA | 83%<br>(20 of 24) | 72-89%*<br>(13-16 of 18) | 85%<br>(35 of 41) | 72-89%*<br>(13-16 of 18) | 83%<br>(20 of 24) | 43-51%*<br>(15-18 of 35) |
| Toxin EIA | 42%<br>(10 of 24) | 100%<br>(18 of 18) | 39%<br>(16 of 41) | 100%<br>(18 of 18) | 42%<br>(10 of 24) | 83%<br>(29 of 35) |

\* GDH EIA results for 3 patients were unable to be retrieved, and all 3 had been categorized as CDI-Unlikely. Clinical specificity was consequently calculated first assuming all 3 GDH EIA results were positive, and second assuming all 3 GDH EIA results were negative.

## DISCUSSION

The high sensitivity of NAAT in diagnosing CDI has been well documented (5). This attribute has been instrumental in directing CDI treatment and avoiding progression of disease for patients with true CDI. However, it is also recognized that molecular tests may lead to overdiagnosis of CDI due to suboptimal clinical specificity and result in increases in both direct and indirect costs related to unnecessary treatment and isolation precautions, overutilization of infection prevention and control resources, and reporting of inflated CDI rates (10-13). Furthermore, CDI treatment with antibiotics has been reported to increase the risk of vancomycin-resistant enterococci (14, 15). Because of this, supplementation of NAAT with Toxin EIA as part of a 2-step testing algorithm has been proposed by IDSA/SHEA and European Society of Clinical Microbiology and Infectious Diseases guidelines (6, 16). Prior studies have reported that performing both NAAT and Toxin EIA may provide clinically useful information. For instance, patients with diarrhea who are NAAT-positive but Toxin EIA-negative may not develop adverse outcomes, even without specific therapy for CDI (17). Also, recurrence of CDI may be more common when both NAAT and toxin assays are positive than when NAAT alone is positive (17). And, complications are more common among patients who are positive by both NAAT and a 3-step algorithm - including GDH, toxins A and B, and cell culture cytotoxicity assay - than when NAAT alone is positive (18).

Nevertheless, others have found that addition of Toxin EIA to NAAT does not reliably distinguish patients with CDI from those who are colonized. Rios et al. found that 75% of their NAAT-positive specimens were Toxin EIA-negative, and that Toxin EIA-negative results were seen just as frequently in CDI as in colonized patients with positive NAAT results (19). We similarly found that most (62%) NAAT-positive specimens were Toxin-EIA-negative, and most (85%) of these patients still benefited symptomatically from CDI treatment. Toxin-EIA results were not contributory to the diagnosis of CDI, because of CDI-Likely patients, Toxin-EIA was negative in 58%. Consistent with this, informal polling (data not shown) of healthcare providers at our institution revealed that virtually all would treat for CDI if it was in the differential diagnosis and the patient had a positive NAAT result, regardless of the Toxin-EIA result.

Because there is currently no gold standard for the diagnosis of CDI, assay performance is particularly challenging to assess. When defining “true positives” and “true negatives” as concordance between both reviewers in categorizing patients as CDI-Likely and CDI-Unlikely, respectively, NAAT performed well (clinical sensitivity 100% and specificity 94%). However, when applying this definition, which uses only the most clinically apparent CDI patients, Toxin EIA had low clinical sensitivity (42%). When including CDI-Indeterminate patients and patients that had discordant categorization into the “true positives”, the clinical sensitivities and specificities of NAAT and Toxin EIA were essentially unchanged; however, when including CDI-Indeterminate/discordant patients as “true negatives”, clinical specificity of NAAT dropped markedly (94% to 49%), and it also decreased for Toxin EIA (100% to 83%). Regardless of how “true” CDI status was defined, the clinical sensitivity of Toxin EIA (range of 39-42%) clearly demonstrated a need for improvement, which was consistent with the findings of others (15, 19, 20).

This study has several limitations. The lack of diagnostic gold standard for CDI resulted in variability between the two reviewers when categorizing patients into the three different CDI status groups. Upon unblinding, the discordant categorizations were thought to be attributable to differences in interpretation of what constituted clinical improvement with treatment for CDI, and lack of agreement as to whether or not the presence of a rectal tube was indicative of a patient having 3 or more loose stools per day. Furthermore, due to the retrospective nature of this study, it was challenging to obtain complete information from the medical records; in particular, frequency of diarrhea, character of stools, and duration of symptoms prior to hospital presentation were difficult to ascertain for some patients. Another limitation of this study was that specimen selection was not completely randomized and was more dependent on when the investigator performing EIA testing was available. Our sample set is likely enriched with NAAT-positive patients; thus, the calculated clinical sensitivities and specificities may not be representative of an unbiased population.

In summary, the diagnosis of CDI remains challenging, but the addition of Toxin EIA testing to NAAT-positive specimens as part of a two-step diagnostic algorithm would provide minimal clinical benefit. At our institution, we have implemented multiple layers of checks to optimize *C. difficile* NAAT utilization, from decision support during ordering to rejection of formed stools during laboratory receipt, and in this setting, NAAT testing alone is currently the best solution for our healthcare system.

## ACKNOWLEDGEMENTS

TECHLAB provided C. DIFF QUIK CHEK COMPLETE tests.

